# Chlorine inactivation of *Escherichia coli O157:H7* in fresh produce wash process: effectiveness and modeling

**DOI:** 10.1101/2021.02.12.430939

**Authors:** Mohammadreza Dehghan Abnavi, Chandrasekhar R. Kothapalli, Daniel Munther, Parthasarathy Srinivasan

## Abstract

This study presents a modified disinfection kinetics model to evaluate the potential effect of organic content on the chlorine inactivation coefficient of *Escherichia coli O157:H7* in fresh produce wash processes. Results show a significant decrease in the bactericidal efficacy of free chlorine (FC) in the presence of organic load compared to its absence. While the chlorine inactivation coefficient of *Escherichia coli O157:H7* is 70.39 ± 3.19 L.mg^−1^.min^−1^ in the absence of organic content, it drops by 73% in chemical oxygen demand (COD) level of 600 - 800 mg.L^−1^. Results also indicate that the initial chlorine concentration and bacterial load have no effect on the chlorine inactivation coefficient. A second-order chemical reaction model for FC decay, which utilizes a percentage of COD as an indicator of organic content in fresh produce wash was employed, yielding an apparent reaction rate of (9.45 ± 0.22) × 10^−4^ μM^−1^.min^−1^. This model was validated by predicting FC concentration (*R*^2^ = 0.96) in multi-run continuous wash cycles with periodic replenishment of chlorine.

## 1. Introduction

Fresh produce and leafy greens play a role in approximately 27% of outbreaks in the United States, according to the Center for Disease Control and Prevention (Chen & Hung, 2017). One critical step in postharvest processing of most fresh produce is the wash with a sanitizing agent to inactivate the microbial load. When the sanitizer concentration drops below a certain level, not only do more pathogens survive, their potential for cross-contamination (i.e., transfer from contaminated to uncontaminated produce) through water also increase. Various types of sanitizers such as free chlorine (Abnavi, Kothapalli, & Srinivasan, 2021; Luo et al., 2012; Tudela, López-Gálvez, Allende, & Gil, 2019), chlorine dioxide (Banach et al., 2017; López-Gálvez, Tudela, Gil, & Allende, 2020), peroxyacetic acid (López-Gálvez, Truchado, Tudela, Gil, & Allende, 2020), activated persulfate (Qi, Wang, Huang, & Hung, 2020), ozone (Gibson, Almeida, Jones, Wright, & Lee, 2019; Hirneisen, Markland, & Kniel, 2011)), gallic acid (Cossu et al., 2016), and sodium acid sulfate (Sheng et al., 2020) have been used for fresh produce sanitization. Despite their disadvantages like the formation of harmful disinfection byproducts, and reduced efficiency in the presence of organic load (Cossu et al., 2016), these sanitizers are relatively cheap, easy to use, and possess high bacterial inactivation efficacy (Teng et al., 2018; Tudela, López-Gálvez, Allende, Hernández, et al., 2019).

Chlorine based sanitizers are widely used in the produce wash industry (Luo et al., 2012), as chlorine is very effective in inactivating pathogens due to its reactive nature (Lee, Yoon, Lee, Han, & Ka, 2010). On the other hand, chlorine also readily reacts with the organic content released from the exudates of fresh produce, and therefore gets depleted (Bolten et al., 2020; Gómez-López, Lannoo, Gil, & Allende, 2014). Meanwhile, excessive levels of chlorine in wash cycles could lead to the formation of undesirable disinfection by-products (DBPs) (López-Gálvez, Tudela, Allende, & Gil, 2019). A critical real-time balance of chlorine levels is thus needed to prevent cross-contamination, as well as minimize the production and accumulation of hazardous DBPs. In this regard, we and others have developed mathematical models which simulate free chlorine (FC) decay in fresh produce wash process (López-Gálvez et al., 2019; Munther, Luo, Wu, Magpantay, & Srinivasan, 2015; Van Haute et al., 2018). Though it is accepted that organic load consumes FC during the wash process, measuring these reactants directly is not simple or straightforward. Studies have linked chlorine decay and physicochemical characteristics of wash water such as chemical oxygen demand (COD), oxidation reduction potential, ultraviolet absorbance at 254 nm (UV254), turbidity, total organic carbon, total suspended solids, and total dissolved solids (Gómez-López, Lannoo, Gil, & Allende, 2014; Li et al., 2019; Van Haute et al., 2018). These studies have afforded successful predictions of the chlorine demand in a single wash process or in pooled process wash water, but not in industry-relevant recycling wash processes with periodic chlorine replenishment. The mechanisms by which chlorine decays in a commercial wash process, and how chlorine loses its efficacy in the presence of organic matter, remain unclear.

Several studies have shown that the efficacy of sanitizers is significantly reduced in the presence of organic matter (Gómez-López et al., 2014; Qi et al., 2020; Teng et al., 2018). However, a model for chlorine inactivation of pathogens in the presence of organic load is not presented yet. Therefore, in this study, we model chlorine decay in the presence of organic matter and investigate the effect of organic load on chlorine inactivation of *Escherichia coli O157:H7* in a model system of iceberg lettuce wash process. We have previously developed a model for chlorine decay in produce wash systems using COD as an indicator for organic load (Munther, Luo, Wu, Magpantay, & Srinivasan, 2015; Srinivasan, Dehghan Abnavi, Sulak, Kothapalli, & Munther, 2020). To investigate how chlorine inactivates pathogens in the presence of organic load, we have presented a disinfection kinetics model. We utilized chlorine to inactivate a three-strain cocktail of non-pathogenic *E. coli O157:H7* and calculated the maximum chlorine inactivation coefficient in the absence of organic load. Then, using our developed model for FC decay and pathogen cross-contamination dynamics, we quantified the chlorine inactivation coefficients in a multi-run wash process of fresh iceberg lettuce. Finally, we demonstrate how chlorine loses its sanitizing efficacy by comparing chlorine inactivation coefficient in the presence and absence of organic content.

## 2. Materials and Methods

### 2.1. *E. coli* O157:H7 culture preparation

Three strains of *E. coli O157:H7* (ATCC-35150, ATCC-43895, and ATCC-1428) were selected for this study. After opening lyophilized vials of each strain, one loop of frozen culture was transferred into Tryptic Soy Broth (TSB) and incubated in a shaking incubator (120 *rpm)* overnight (37 °C). The incubated broth was further sub-cultured in TSB with nalidixic acid until the final broth had 50 mg/L nalidixic acid. After incubation, cells were harvested by centrifuging at 3000 × *g* for 10 min, and the collected cells were washed twice with sterile phosphate buffered saline (1× PBS) and subsequently resuspended in 50 mL of PBS. Equal volumes of each strain were mixed to make the final *E. coli* cocktail with approximately 9-log MPN.mL^−1^. This bacterial cocktail was used to prepare 5- and 6- log MPN.mL^−1^ solutions for disinfection experiments.

### 2.2. Lettuce preparation and inoculation

Iceberg lettuce was purchased from local grocery stores and used on the same day of purchase for experiments. After removing outer leaves, fresh leaves were chopped into 1”×1” pieces and stored at 4 °C until the start of experiments. Separately, red leaf lettuce was purchased to be inoculated and added to the washing flume as inoculated produce. After removing outer leaves and chopping fresh red leaf lettuce leaves into 2”×2” pieces, 20 droplets of 5 μL *E. coli* cocktail with 6 log MPN.mL^−1^ were placed per gram of chopped red leaf lettuce to obtain a final *E. coli* concentration of 5-log MPN.g^−1^. Inoculated red leaf lettuce were kept refrigerated (4 °C) overnight to let the bacteria attach to the surface of leaves.

### 2.3. Inactivation of *E. coli O157:H7* by chlorine

These experiments were designed to determine chlorine inactivation (lethality) coefficient of *E. coli O157:H7* in the absence of organic load. All disinfection experiments were done in 500 mL flasks containing 250 mL of tap water. The pH was regulated to 6.5 by adding 0.1 M citric acid. After sterilizing flasks for 20 min at 121 °C, an appropriate density of cells from the *E. coli* cocktail were transferred to the flasks to yield a final *E. coli* concentration at 5- and 6- log MPN.mL^−1^, and the flasks were refrigerated (4 °C). Then 0.7, 1.4, or 2.8 mL of 1000-fold diluted 4.5% sodium hypochlorite (BCS Chemicals, Redwood City, CA, USA) was added to the flasks to achieve 0.125, 0.25, or 0.50 mg.L^−1^ initial free chlorine (FC) concentration solutions. Flasks were continuously mixed (200 *rpm)* using an overhead stirring apparatus equipped with sterile paddles. Samples (1 mL) were taken from the reaction vessels at the desired contact times and added to tubes containing 9 mL deionized water with 0.1% (wt./vol.) sodium thiosulfate (Sigma Aldrich) to immediately neutralize residual chlorine. Chlorine concentrations were determined immediately after taking the sample, using the N, N-diethyl-p-phenylenediamine (DPD) method, with a Chlorine Photometer (CP-15, HF Scientific Inc., Ft. Myers, FL). Bacteria survival was measured by counting cells via modified Most-Probable-Number (MPN) method using 48-well deep microplates (Luo et al., 2012). All experiments were independently replicated three times.

### 2.4. Chlorine decay in single-wash (batch) experiments

The goal of these experiments was to model FC decay due to the reaction with organic matter released from produce exudate. Concentrated (4.5%) sodium hypochlorite was added to 3 L of tap water (maintained at 4 °C) to attain FC concentration ~26 mg.L^−1^ (~498 μM). The pH was adjusted to 6.5 with 0.1 M citric acid. Before starting the experiment, samples were collected to measure FC and COD. Within 5 min of adding chlorine, 250 g chopped iceberg lettuce was washed in that water for 30 seconds and removed by a sieve. Aliquots to measure FC and COD were taken every two minutes after washing for the next 30 minutes.

### 2.5. *E. coli* O157:H7 inactivation in continuous lettuce wash process

These experiments were designed to investigate whether the chlorine inactivation coefficient of *E. coli* remains similar even in the presence of organic matter. Accordingly, these studies were done in a large tank containing 20 L of tap water maintained at 4 °C. Chopped iceberg lettuce with inoculated red leaf lettuce pieces entered the tank from one side and exit from the other side. Before adding chopped lettuce, concentrated (4.5%) sodium hypochlorite was added to the tank to achieve initial FC levels ~12 mg.L^−1^ (~230 μM). The pH was adjusted to 6.5 by adding 0.1 M citric acid. Within 5 min of adding chlorine, chopped iceberg lettuce was continuously introduced to the tank at a rate of 0.5 kg.min^−1^, along with inoculated red leaf lettuce at the rate of 10 g.min^−1^. The dwell time of the lettuce pieces was set to 30 sec, and each run lasted ten minutes.

Samples were taken every two minutes after starting the experiment to measure FC, COD, and bacterial load in the wash water *(X_W_)* and on the surface of iceberg lettuce *(X_L_)*. COD was measured using the reactor digestion method (Jirka & Carter, 1975). Wash water samples for bacterial analysis were treated with 0.1% (wt./vol) sodium thiosulfate to rapidly quench residual chlorine. To measure *E. coli* amounts transferred to the surface of uninoculated iceberg lettuce, two 25-g samples of iceberg lettuce slices exiting from wash tank were weighed and stored in sterile filter bags (WhirlPak®, Nasco, Modesto, CA) and incubated with 125 mL of sterile tryptic soy broth (TSB) at 230 *rpm* for 2 min with a stomacher blender (Seward Stomacher 400, London, UK). The *E. coli* levels in the filtrate or water was enumerated using the modified MPN method (Lou et al., 2012).

After ten minutes and washing of 5 kg of iceberg lettuce, sodium hypochlorite was added to the tank to replenish chlorine and pH was adjusted to 6.5 using 0.1 M citric acid. The second run was started five minutes after the end of first run. Similarly, the third run was performed using similar procedures as the first and second runs. A total of 15 kg of iceberg lettuce were washed by the end of these three runs, and this experiment was independently repeated three times.

### 2.6. Chlorine decay and bacterial disinfection model in produce wash processes

We have previously reported that chlorine concentration could be predicted using a percentage of COD level change in wash water, which also serves as an indicator for organic load (Srinivasan, Dehghan Abnavi, Sulak, Kothapalli, & Munther, 2020):

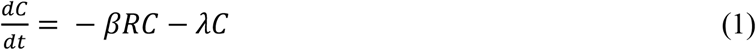

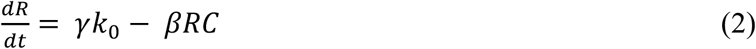

where *k_0_* (μM.min^−1^) stands for the rate of addition of organic matter to the washing flumes measured by the rate of change in COD levels. Namely,

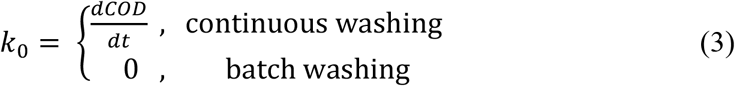

Equation 1 depicts the decay rate of chlorine with the FC levels at any time *t* given by *C* (μM), the concentration of organic load that reacts with chlorine given by *R* (μM), and the apparent reaction rate constant for the reaction of FC and organic matter in wash water given by *β* (μM^−1^.min^−1^). As the reactant concentration *R* introduced to the wash water by produce is not measurable directly, it has been replaced by a fixed percentage (γ) of COD change. It should be noted here that organic load is not proportional to the COD level, but to the change in COD level. Finally, *λ* (min^−1^) denotes the first-order natural decay rate of chlorine. The second term on the right-hand side of equation 2 models the consumption rate of organic load by FC as a second-order reaction. The first term on the right-hand side of this equation is the rate at which the organic load is added to the washing flume. The initial conditions for equations 1 and 2 are *C*(0) = *C*_0_ and *R*(0) = γ(ΔCOD), where *C_0_* (μM) is the initial chlorine concentration and *ΔCOD* (μM) indicates the COD change (before and after wash) for a single wash experiment. We have used the fixed percentage of COD change before and after wash as an initial concentration of organic matter added to the wash vessel.

The concentration of pathogens in the process water, *X_W_* (MPN.mL^−1^), could be described as (Munther, Luo, Wu, Magpantay, & Srinivasan, 2015):

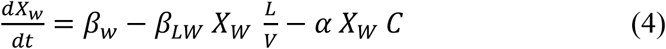

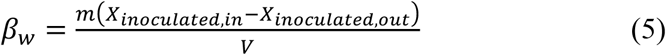

where *β_w_* (MPN.mL^−1^.min^−1^) is the shedding rate of *E. coli* to washing system, *m* (g.min^−1^) is the adding rate of inoculated produce, *V*(mL) is the tank volume, *X_inocuiated,in_* and *X_inocuiated,out_* are the respective concentrations of *E. coli* on the surface of inoculated produce entering and exiting the wash vessel. The second and third terms in equation 4 account for the removal mechanisms of pathogens from the wash tank. The second term of equation 4 represents the rate of binding of *E. coli* to uninoculated produce. This binding rate depends on the concentration of pathogens in the wash water (*X_W_*) and the ratio of uninoculated produce in the wash tank to the wash tank volume (L/V (g/mL)), with *β_LW_* (mL.g^−1^.min^−1^) being the proportionality constant. We model the inactivation of suspended pathogens by FC *via* the third term of equation 4 where *C* (μM) is the concentration of FC and *α* is the inactivation (lethality) coefficient of *E. coli* by FC (μM ^−1^.min^−1^).

The contamination dynamics for the uninoculated produce depends on the binding rate *(i.e.* the rate at which pathogens in the water bind to lettuce), the FC inactivation of *E. coli,* as well as the average time lettuce spends in the wash tank (Munther, Luo, Wu, Magpantay, & Srinivasan, 2015):

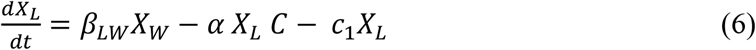

where *X_L_* (MPN.g^−1^) is the pathogens level on the lettuce surface. The first term in equation 6 indicates the rate of *E. coli* transfer from water to lettuce, while the second term reflects the inactivation of pathogens on lettuce by FC. For the third term, we assume that the exit time of lettuce from the wash tank is exponentially-distributed whose mean is 1/*c1*, *i*.*e*., 1/*c1* (min) reflects the average dwell time for lettuce in the wash tank. It should be noted that we did not include produce to produce transmission of the pathogens in our model.

Equation 4 shows the bacterial concentration in the water in continuous wash of produce. In case of no produce wash (experiments of section 2.3), the first two terms are zero, and only the last term remains. In this case, the model reduces to the Chick-Watson disinfection model (Watson, 1908). Previous studies (Banach et al., 2017; Qi et al., 2020; Teng et al., 2018) have shown that organic load impacts sanitization by depleting as well as reducing the efficacy of the sanitizer. So, in order to consider the effect of organic load on the bactericidal efficacy of chlorine, the inactivation coefficient *α* is modeled as:

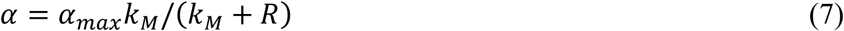

Here, *α_max_* (μM^−1^.min^−1^) is the maximum inactivation coefficient in the absence of organic load, *R* (μM) is the concentration of the organic reactants, and *k_M_* (μM) is a constant. Analogous to Michaelis-Menten kinetics, the value of *k_M_* is numerically equal to the concentration of organic reactants at which the inactivation rate is half of *α_max_*. Equation 7 shows that when the concentration of organic load rises in the wash process, the sanitizer efficacy decreases. To account for this dependence of *α* on *R,* we use its form from equation 7 in equations 4 and 6.

### 2.7. Parameters determination and statistical analysis

All experiments were carried out in triplicate. MINITAB Statistical Software package (Version 17) was used to perform one-way analysis of variance (ANOVA) and Tukey’s test. A *p*-value < 0.05 between groups was considered statistically significant. Three Python packages - *statsmodels, SciPy* and *lmfit* were used to optimize model parameters.

#### 2.7.1. Maximum inactivation coefficient determination

Maximum inactivation coefficient (*α_max_*) of *E. coli* by FC was obtained from inactivation experiments data with no organic load (section 2.3) and equation 4. For these experiments, there is no produce wash, and so the first two terms of equation 4 as well as the value of *R* equal to zero. So, equation 4 (replacing *X_w_* with *N)* for this case will be:

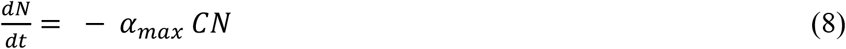

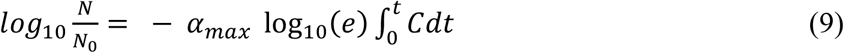

where *N* (MPN.mL^−1^) is the density of surviving bacteria after time *t* (min), *N_0_* (MPN.mL^−1^) is the initial concentration of bacteria, *C* (mg.L^−1^) is the chlorine concentration, and α_max_ is the inactivation coefficient (L.mg^−1^.min^−1^) in the absence of organic matter. Equation 8 is Chick-Watson disinfection model. In order to be consistent with the literature, we have used mg.L^−1^ as the unit for the chlorine concentration (*C*), so that the unit for *α* is L.mg^−1^.min^−1^. Others have reported the value *α_max_* log_10_(e) ≈ 0.4343 *α_max_* as the inactivation coefficient (Lee, Yoon, Lee, Han, & Ka, 2010; Cho et al., 2010).

FC decays over time due to the reaction with organic matter. Defining *CT* value (mg.min.L^−1^) as 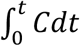 and given by the area under the chlorine decay curve (**Fig. 1**), we calculated *CT* for all experiments by applying Simpson’s Rule.

**Figure 1.**
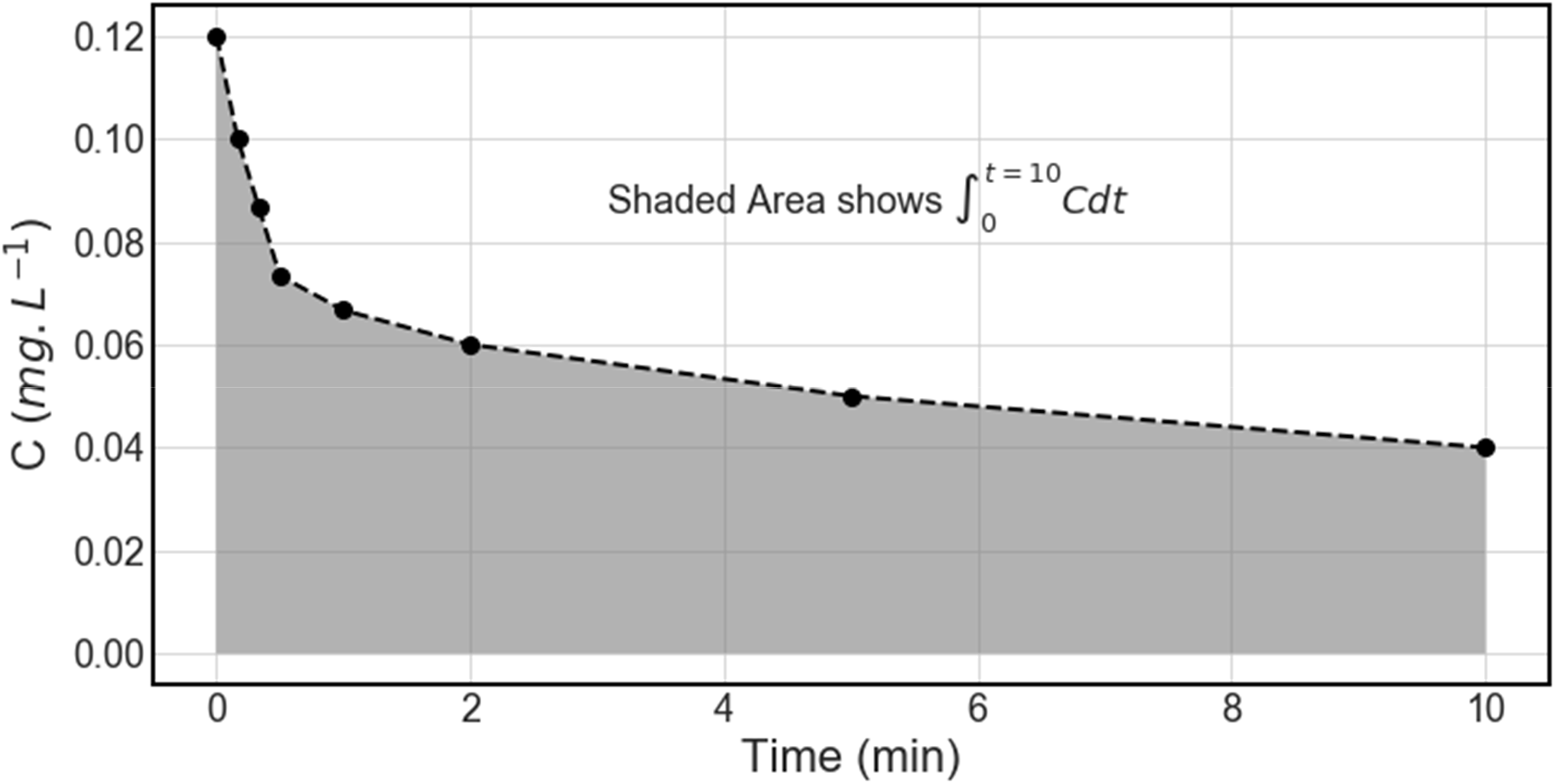
Schematic representation of *CT* value when initial FC concentration was 0.12 mg.L^−1^. The shaded area shows the *CT* value (the integration part of equation 9) for ten minutes after adding chlorine to the solution.

The left-hand side of equation 9 is zero at *t* = 0 (*i.e*., when *N*=*N_0_*) and approaches a minimum when the surviving bacteria becomes undetectable. Assuming *t*^*^ as the time it takes for the bacterial concentration to become undetectable, equation 9 was modified as:

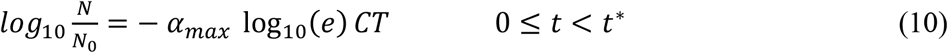

In our study, the lowest bacterial detection level is 0.12 MPN.mL^−1^.

#### 2.7.2. Apparent reaction rate constant determination

To find the optimal values for *γ* and *β,* we used the results from batch experiments (section 2.4). To begin with, the decay rate of FC in water *(λ)* is set to 1.7 *×* 10^−3^ min^−1^ (Srinivasan, Dehghan Abnavi, Sulak, Kothapalli, & Munther, 2020). Since the stoichiometric coefficients of the reaction between FC and organic compounds (FC + *R* → *Products*) are similar for both FC and organic matter (Deborde & von Gunten, 2008), the change in the concentrations of FC and organic reactants should be similar, i.e., R_0_ — R = *C_0_* — *C.* At the end of the batch experiment when there is no change in FC, we assume that all organic matter has been consumed by FC and *R_end_* ≅ 0. Having this point and replacing R_0_ = *γ(ΔCOD),* we obtain

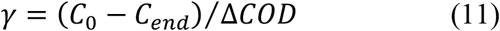

This equation for *γ* could thus be used to find the organic load at each time point:

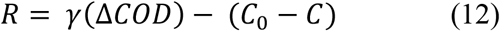

Using the experimental data for FC, estimated organic content (*R*) from batch experiments, and the model described in equations 1-2, we obtained *β* via curve-fitting using the Levenberg-Marquardt algorithm.

#### 2.7.3. Binding rate constant and inactivation coefficient determination in continuous wash

After determining *γ* and *β* we validated our model by predicting the FC concentration in continuous experiments and comparing our results with experimental data. Finally, with the bacteria survival data from continuous experiments (section 2.5) and using the model explained in equations 4-7, we obtained the optimal values for *k_M_* and *β_LW_* with Levenberg-Marquardt algorithm.

#### 3. Results and Discussion

##### 3.1. *E. coli* inactivation by free chlorine

The time-dependent drop in FC and *E. coli* levels (pH ~ 6.5; 4 °C) at 5 or 6 MPN.mL^−1^ of initial *E. coli* concentration and *C_0_* from 0.12 - 0.5 mg.L^−1^ was shown in **Fig. 2**. The FC decay was higher in runs with 6-log MPN.mL^−1^ *E. coli* within the first 2 minutes. Similar trends were noted at other conditions, and the change in FC levels was insignificant after the first two minutes. At *C_0_* ≥ 0.25 mg.L^−1^, all bacteria were inactivated after the first two minutes. At *C_0_* = 0.12 mg.L^−1^, total inactivation of bacteria needed more time due to low FC concentration. Expectedly, these results also suggest that the higher the *C_0_*, the faster the inactivation of *E. coli*. For example, a 3-log reduction in *E. coli* (99.9% inactivation) was achieved after 10 seconds when *C_0_* = 0.5 mg.L^−1^, while it took 30 seconds when *C_0_* = 0.25 mg.L^−1^ and even longer when *C_0_* = 0.12 mg.L^−1^. It took more than a minute to achieve 3-log reduction in *E. coli* levels when *N_0_* = 5-log MPN.mL^−1^, and two minutes when *N_0_* = 6-log MPN.mL^−1^.

**Figure 2.**
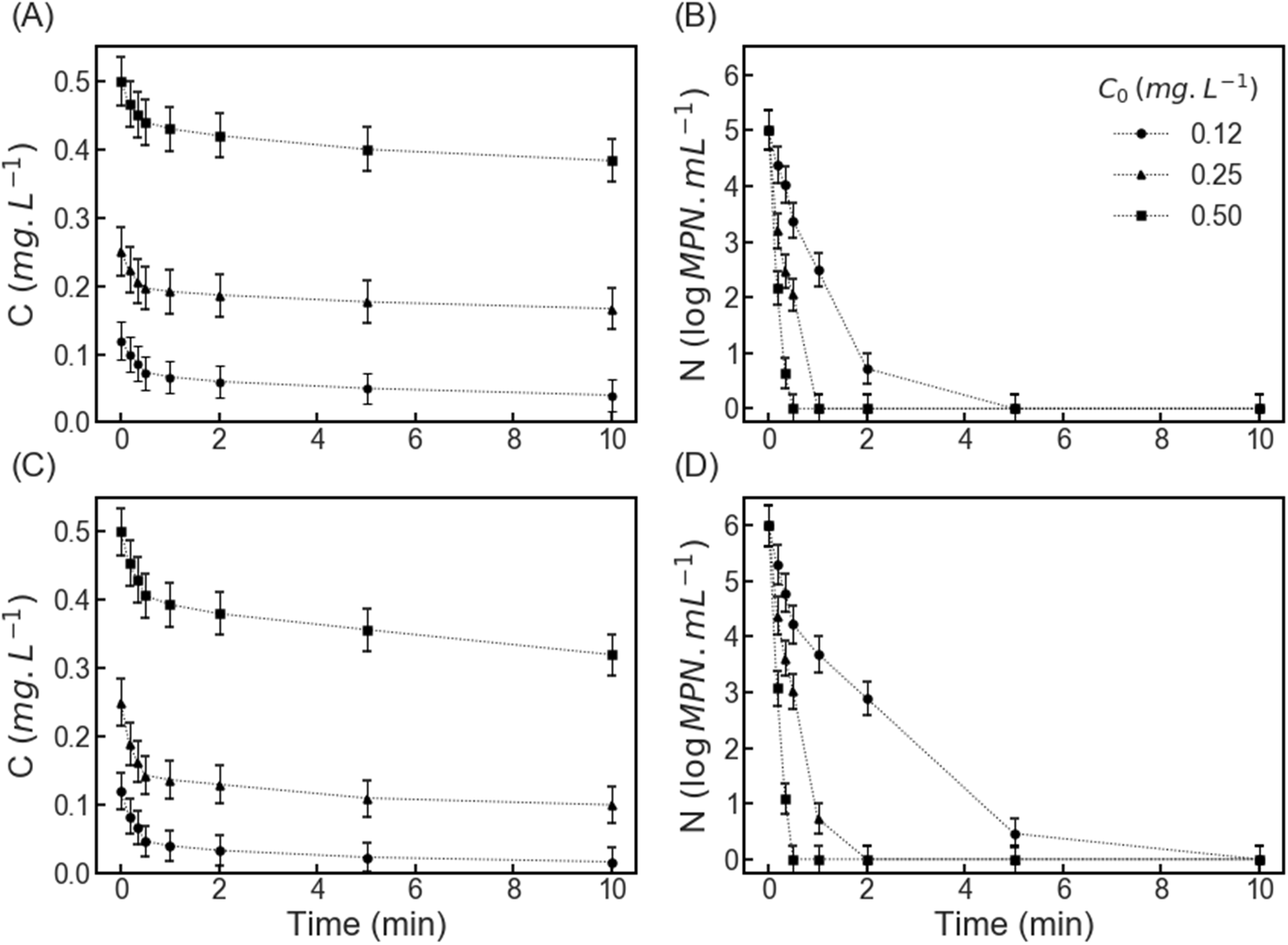
Free chlorine decay in the presence of initial *E. coli* levels at (A) 5-log MPN.mL^−1^ and (C) 6-log MPN.mL^−1^, at pH ~ 6.5 and wash water temperature at 4 °C. Inactivation of (B) 5-log MPN.mL^−1^ and (D) 6-log MPN.mL^−1^ of *E. coli* at three concentrations of FC: 0.12, 0.25 and 0.5 mg.L^−1^. The symbols are arithmetic averages of three independent experiments and the error bars indicate respective standard deviations. The dotted lines connecting symbols were for visual aid.

The time-dependent *E. coli* reduction within the first minute of contact with FC was shown in **Fig. 3**. Regardless of initial bacterial load (*N_0_*), if the chlorine level is enough, the *E. coli* reduction ratio is the same for all starting FC levels (*C_0_*). Thus, it can be conjectured that *E. coli* inactivation coefficient is not a function of initial bacterial load and FC concentration. Equation 10 which explains the disinfection kinetics in the absence organic content (section 2.7.1) was used to calculate the maximum inactivation coefficients *(α_max_*) at various *C_0_* and *N_0_*, and in turn, the effects of *C_0_* and *N_0_* on maximum inactivation coefficient was investigated using ANOVA. ANOVA analysis (Supplementary **Table S1**) showed that inactivation coefficient was indeed independent of the initial chlorine concentration (*C_0_*) and bacterial load *(N_0_)*.

**Figure 3.**
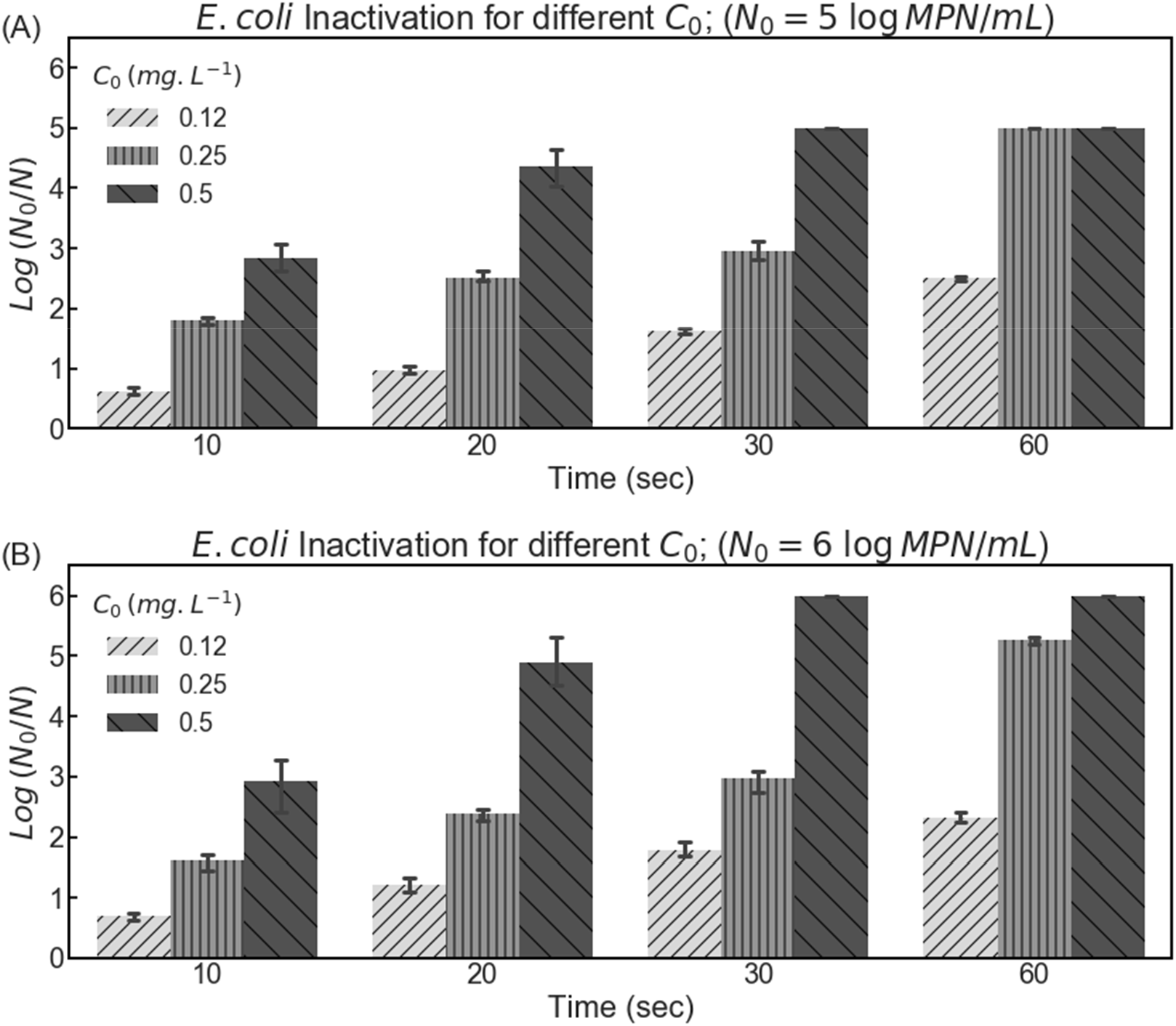
*E. coli* reduction by free chlorine at various initial FC concentrations (*C_0_*: 0.12, 0.25, and 0.5 mg.L^−1^) within the first one-minute of contact time (pH ~ 6.5 and 4 °C). Error bars show standard deviation.

The first step towards finding the inactivation coefficient is to calculate CT values for each experiment. CT value is the product of contact time and chlorine concentration which can be determined by integrating the chlorine decay profile (**Fig. 2 - A, C**). After obtaining CT values and results from bacterial disinfection experiments (**Fig. 2 - B, D)**, the inactivation coefficients were calculated using equation 10 and tabulated (**Table 1**). The maximum inactivation coefficient *(α_max_*) we attained was similar to that reported by others (Lee, Yoon, Lee, Han, & Ka, 2010) using the Chick-Watson model, at 4 °C and pH ~ 8.5.

**Table 1.** Mean values of *E. coli* inactivation (lethality) coefficient *(α_max_*) by FC under different initial FC level and *E. coli* concentration. Errors show 95% confidence interval.

| $N_0$<br>(log MPN. mL<sup>-1</sup>) | $C_0$<br>(mg. L<sup>-1</sup>) | $\alpha_{max}$<br>(L.mg<sup>-1</sup>.min<sup>-1</sup>) | $R^2$ |
| --- | --- | --- | --- |
| 5 | 0.12 | 70.05 ± 3.65 | 0.99 |
|  | 0.25 | 70.49 ± 22.68 | 0.97 |
|  | 0.50 | 67.74 ± 20.89 | 0.98 |
| 6 | 0.12 | 76.73 ± 8.27 | 0.99 |
|  | 0.25 | 78.38 ± 7.07 | 0.99 |
|  | 0.50 | 76.35 ± 13.28 | 0.99 |

Results presented in **Figures 4 and 5** suggest that the disinfection model, presented in equation 10, describes chlorine inactivation of *E. coli* very well. It is evident that there is no significant difference in the slopes of lines (*i*.*e*., inactivation coefficients) at varying initial levels *C_0_* (**Fig. 4**). Similarly, bacteria initial load has no effect on their inactivation coefficient by chlorine (**Fig. 5**). The average inactivation coefficient *(α_max_)* was 70.39 ± 3.19 L.mg^−1^.min^−1^ (≈ 30.57 L.mg^−1^.min^−1^/log_10_(*e*) ≈ 3.69 μM^−1^.min^−1^). The CT values for 2- to 4- log inactivation of *E. coli* was 0.035 - 0.125 mg.min.L^−1^. The CT values we noted for inactivation of *E. coli* were approximately 1000 times lower than the corresponding CT values for inactivation of Giardia cyst by free chlorine presented in the surface water treatment rule (SWTR) of USA (USEPA, 1991). Taylor *et al.* (Taylor, Falkinham, Norton, & LeChevallier, 2000) reported CT values in the range of 0.034 - 0.05 mg.min.L^−1^ for a 99% (2-log) reduction of *E. coli* by chlorine, while Helbing et al. (Helbing & VanBriesen, 2007) noted the chlorine contact time for 3-log inactivation of *E. coli* as 0.032 ± 0.009 mg.min.L^−1^, similar to our observations in the current study.

**Figure 4.**
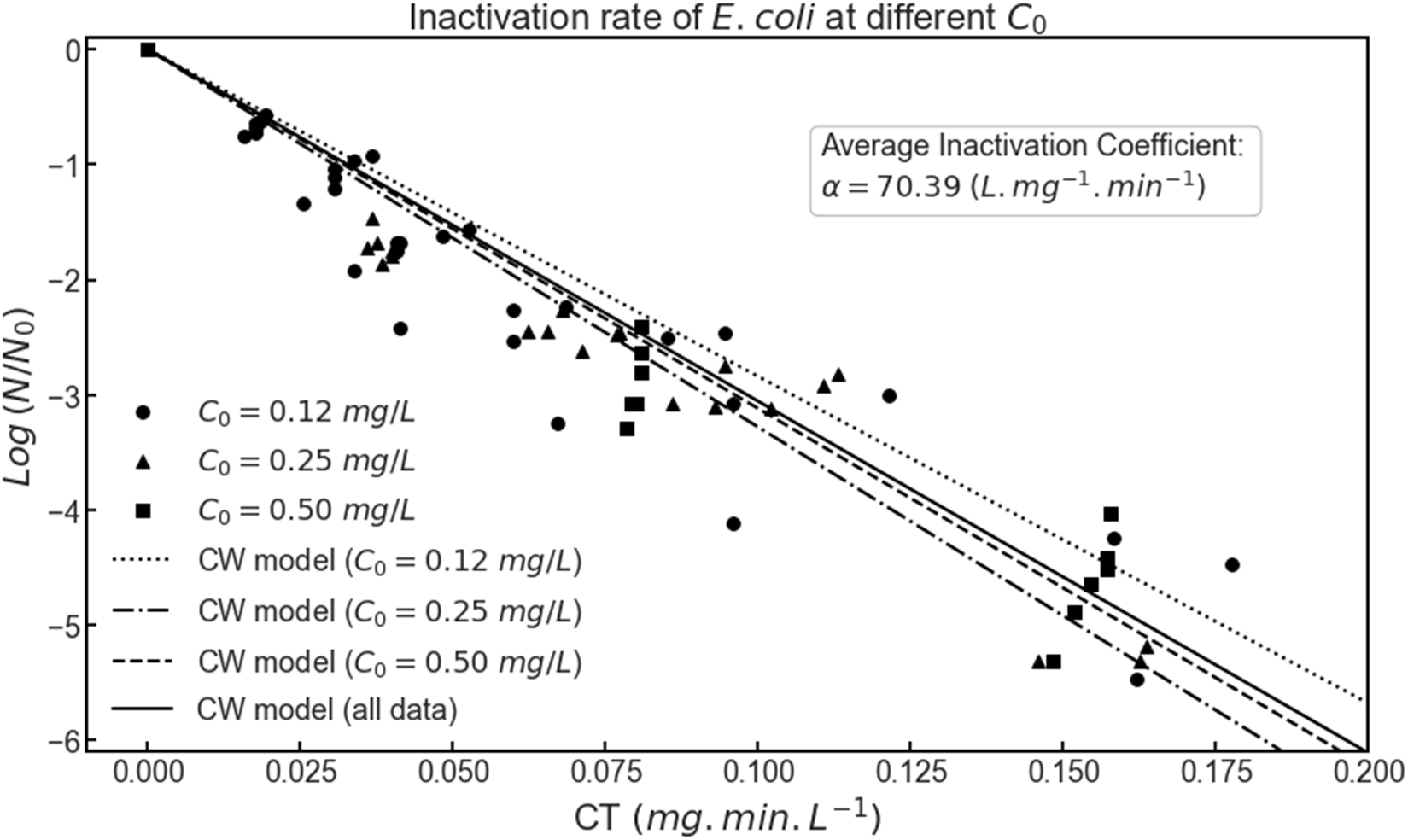
Inactivation of *E. coli* by free chlorine in the absence of organic load at different initial FC levels (0.12, 0.25 and 0.5 mg.L^−1^). The symbols depict experimental data, and the lines represent the fits from the disinfection model (Eq. 8) for *E. coli* inactivation by free chlorine at different FC levels (pH 6.5 and 4 °C).

**Figure 5.**
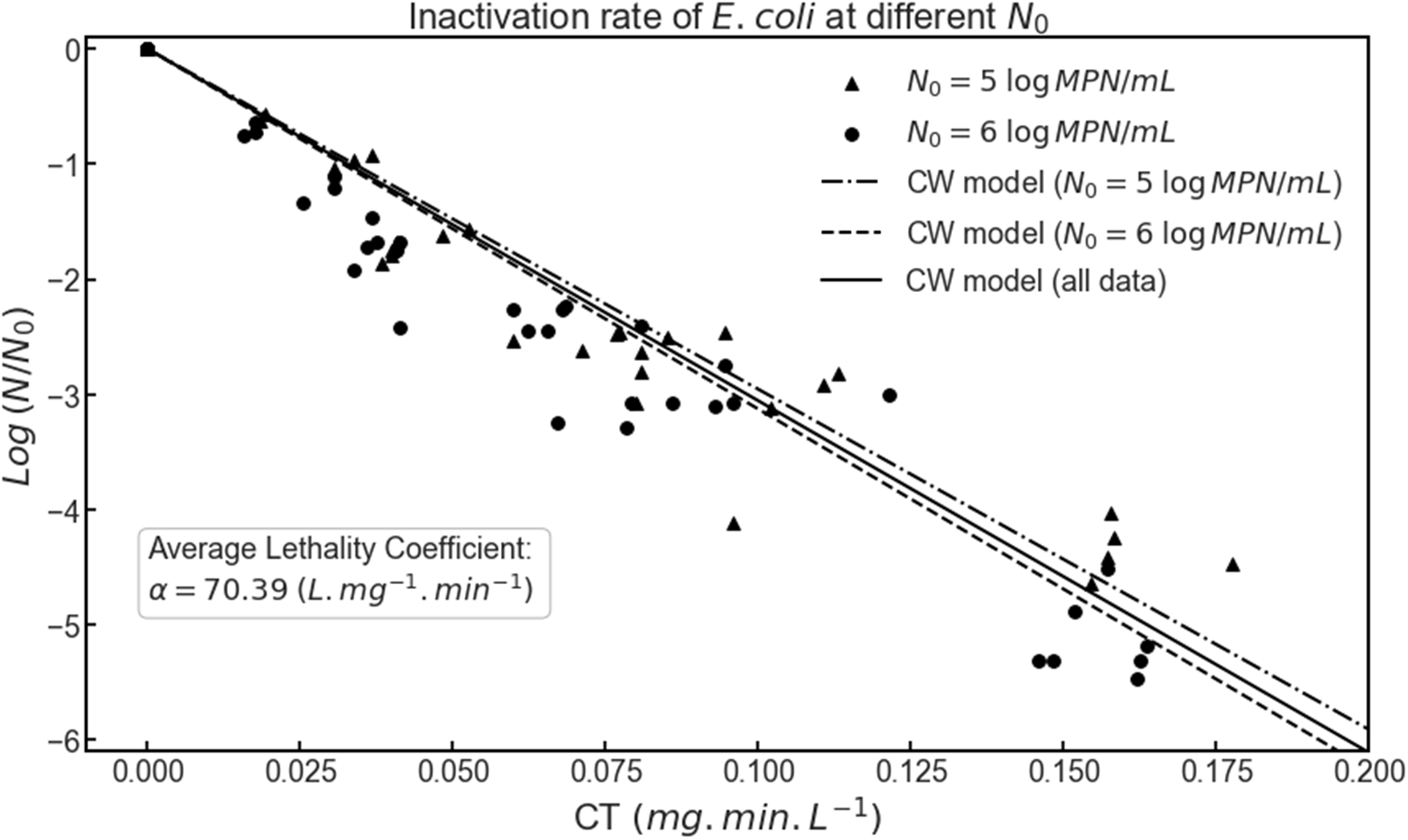
Chlorine inactivation of *E. coli* in the absence of organic load with starting concentrations of 5 and 6 MPN.mL^−1^. The symbols represent experimental data while the lines are fits from the disinfection model (Eq. 8) for *E. coli* inactivation by free chlorine (pH ~ 6.5, 4 °C).

##### 3.2. Establishing chlorine decay model using single-wash experiments

The changes in FC, organic matter, and COD levels in single-wash experiments (section 2.4) were shown in **Fig. 6**. A sharp drop (55%) at the beginning of experiment (first 10 min) was followed by a slow decay of FC (Fig. 6A). Since measuring organic content in real-time is not possible, we used COD levels from wash water as a proxy indicator. The drop in FC concentration was ~ 296 μM after 30 min wash, and while the reaction between FC and organic matter most likely ended after the first twenty minutes. Assuming that the chlorination of most organic load released from produce into the water has the same stochiometric coefficient as FC (Deborde & von Gunten, 2008), the change in organic load should reflect that in FC. Taking the changes in FC and COD levels into account and using equations 11 and 12, *γ* was determined as 0.12 ± 0.01 for iceberg lettuce (**Table 2**). Recall that *γ* is the percentage of ΔCOD and represents the ratio of the change in FC to the change in COD level. While the COD level remained constant over the duration (30 min) of the experiment (horizontal line in Fig. 6B), the concentration of the organic load (*R*) will drop over the course of the experiment. This is the reason why the COD was not directly used in our model.

**Figure 6.**
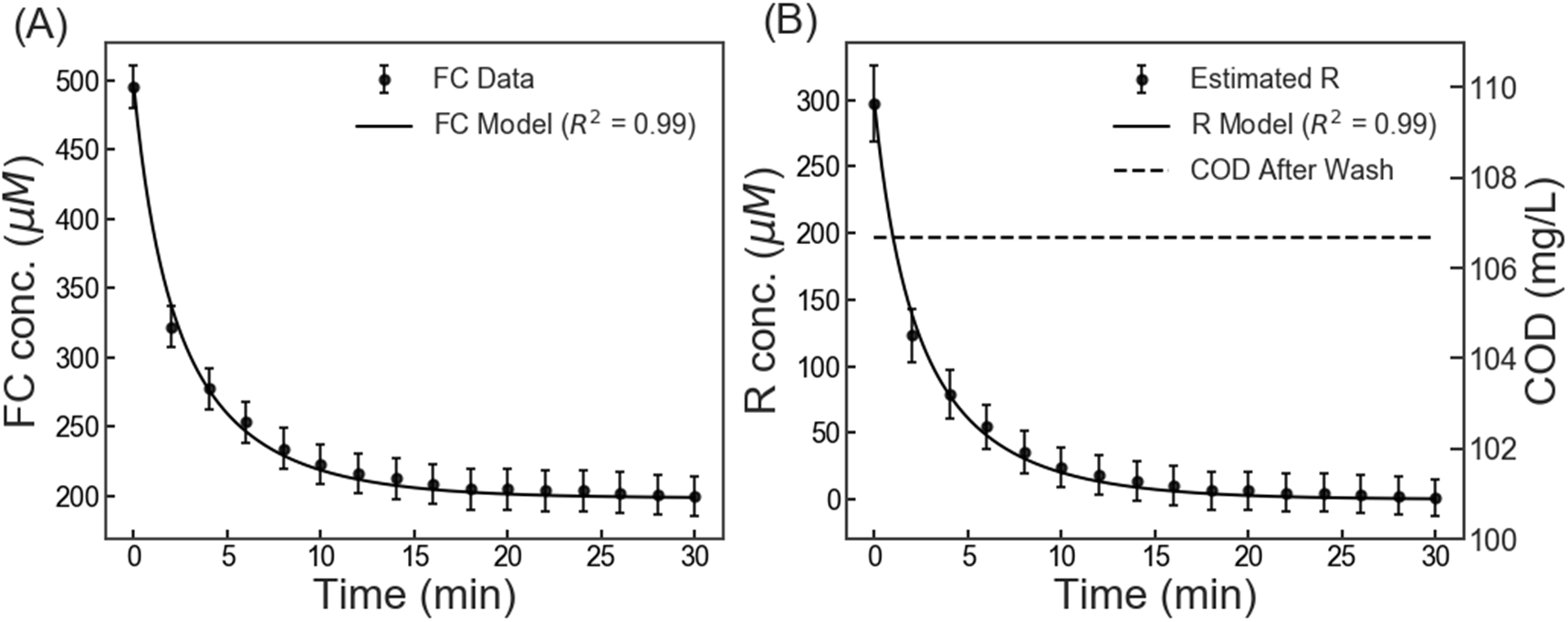
(A) Free chlorine (FC) decay, and (B) organic load consumption and corresponding COD levels obtained from single wash (batch) experiments of 250 g chopped iceberg lettuce. Data (filled circles) was reported as average + standard deviation. The COD levels were shown as the dotted line in (B). The model was trained based on the experimental data to obtain the reaction rate constant, *β* (reported in **Table 2**). *R*^2^ ≥ 0.99 proves that the model fits the experimental data well.

**Table 2.** Results from curve-fitting and model training from single-wash experiments. Errors show the 95% confidence intervals.

| $COD_0^a$<br>($\text{mg.L}^{-1}$) | $COD_{end}^b$<br>($\text{mg.L}^{-1}$) | $\Delta\text{COD}$<br>($\text{mg.L}^{-1}$) | $\Delta\text{COD}$<br>($\mu\text{M}$) | $\Delta\text{FC}$<br>($\mu\text{M}$) | $\gamma$ | $\beta$<br>($\mu\text{M}^{-1} \cdot \text{min}^{-1}$) |
| --- | --- | --- | --- | --- | --- | --- |
| $28.0 \pm 6.7$ | $106.7 \pm 5.8$ | $78.7 \pm 4.1$ | $2458.3 \pm 129.3$ | $296.1 \pm 10.0$ | $0.12 \pm 0.01$ | $(9.45 \pm 0.22) \times 10^{-4}$ |

<sup>a</sup>COD before wash; <sup>b</sup>COD after wash

After calculating *γ* and estimating the organic content at each time point, equations 1 and 2 were fitted to the single-wash experimental data (section 2.4) and the apparent rate constant *(β)* for chlorination of organic load in iceberg lettuce wash process was determined using Levenberg-Marquardt algorithm (**Table 2**). Based on the units used for FC concentration (μM) and time (min), *β* was obtained as 9.45 × 10^−4^ μM^−1^.min^−1^ (≈15.76 M^−1^.s^−1^). The rate constant for the reaction of hypochlorous acid with different types of organic load that could possibly enter the wash water from produce exudate ranged from 10^−2^ M^−1^.s^−1^ to 10^7^ M^−1^.s^−1^ (Deborde & von Gunten, 2008). These organic compounds include amino acids, amides, and phenolic compounds, among others. The advantage of this model is that measuring each of these organic reactants involves complicated analytic methods while measuring COD is relatively quick and straightforward. Moreover, we and other researchers have reported the strong correlation between produce-water ratio and COD levels for various types of produce (Abnavi, Alradaan, Munther, Kothapalli, & Srinivasan, 2019; Li et al., 2019). Therefore, with careful calibration of commercial systems, it is possible that the measurement of COD is redundant.

##### 3.3. Chlorine decay prediction for continuous-wash process

Thus far, we presented a model for chlorine decay (equations 1-2), fitted that to the single-wash experiment data (Fig. 6), and optimized the model parameters (Table 2). To validate the accuracy of our model for large-scale experiments and various experimental conditions, we used our model to predict FC concentration in continuous wash of iceberg lettuce in a 20 L washing flume (section 2.5). These experiments consisted of 5 steps; three 10-min runs of washing iceberg lettuce and two 5-min stops to replenish chlorine and regulate pH. The decay rate of chlorine increased in each successive run due to the accumulation of organic compounds in the washing flume (**Fig. 7A**), which was supported by COD data. The COD level increases constantly over the three wash cycles, with a small dip in COD levels at the two 5-min intervals when chlorine was replenished (**Fig. 7B**).

**Figure 7.**
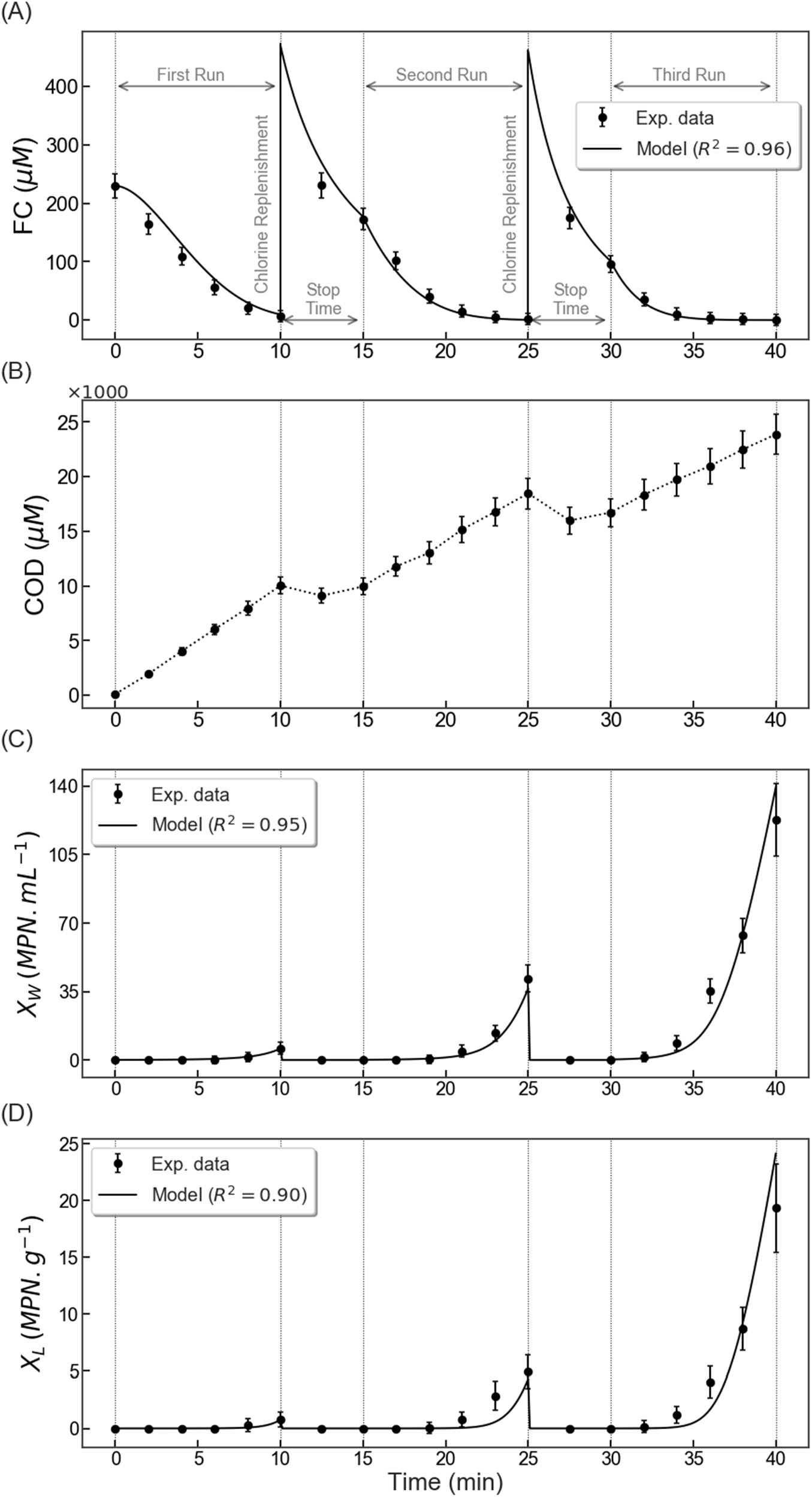
Free chlorine profile (A), COD (B), *E. coli* concentration in water (C), and *E. coli* concentration on iceberg lettuce post-wash (D). Solid circles represent the averages of experimental data with standard deviation, while solid lines are model predictions. Using experimental data from (C) and (D), the model parameters were fitted as *k_M_* = 524 μ_lw_ and *β_w_* = 0.41 mL.g^−1^.min^−1^. The average inactivation coefficients for the three runs of the experiment were 45.05, 28.86, and 19.01 L.mg^−1^.min^−1^ respectively.

We further tested the performance of our model by predicting FC levels in continuous-wash experiments using equations 1-3. The fixed parameters of the model were determined in the previous section using the experimental data from single-wash experiment *(γ* = 0.12; *β* = 9.45 × 10^−4^ μM^−1^ min^−1^). The other required parameter is the rate of organic load entering the wash water (*k_0_*) which could be determined from equation 3 and obtained *via* COD data presented in **Fig. 7B**. Our results (**Fig. 7A**) show that the model predictions for FC match the experimental data very well (*R*^2^ = 0.96). While our model was trained with batch-wise experiments, it was able to accurately predict the FC levels for a continuous wash experiment, suggesting that the model can capture the underlying mechanism of free chlorine decay under different conditions.

##### 3.4. *E. coli* inactivation in continuous wash: Modeling and parameter tuning

The bacterial concentration in the water *(X_W_)* and on the surface of uninoculated produce *(X_L_)* was shown in **Fig. 7** (C, D). The bacterial load increases in successive runs due to the increase in organic load. While FC levels drop to near zero only at the end of the cycle in the first run, the chlorine levels depleted sooner in the second and third runs, leaving a greater chance for *E. coli* survival. Therefore, the *E. coli* levels rose to ~122 MPN.mL^−1^ in water by the end of the third run, while their levels were ~ 6 MPN.mL^−1^ at the end of the first run. As noted in **Fig. 7**, as the COD level increases, FC decays faster, leading to rise of bacterial load in the water, and thus more bacterial transfer to iceberg lettuce.

Earlier (section 3.1), we determined the maximum chlorine inactivation coefficient of *E. coli* as ~70.39 L.mg^−1^.min^−1^ in the absence of organic load. We hypothesized that the organic load would affect the inactivation coefficient and chlorine will lose its sanitizing efficacy in the presence of organic load (equation 7). The inactivation coefficient could be determined from the continuous wash experimental data and the model explained for bacterial survival in produce wash systems (equations 4-6). However, to fit the model to the experimental data, we first need to measure the shed rate of *E. coli* to the system (*β_w_*). As described in section 2.5, the inoculated produce (*m*) was added at a rate of 10 g.min^−1^ with 5-log MPN.g^−1^ *E. coli* concentration. Results show a 1-log reduction in *E. coli* from collected inoculated produce after the wash. Taking the volume of the wash water as 20 L, and using equation 5, *β_w_* was obtained as 45 ± 1.8 MPN.mL^−1^. min^−1^. Finally, the model was fitted to the experimental data and the *E. coli* binding rate to iceberg lettuce *(β_LW_)* and constant *k_M_* were determined to be 0.41 ± 0.07 mL.g^−1^.min^−1^ and 524 ± 114 μM, respectively (**Table 3**). The average *α* for the continuous wash experiments was ~ 64%, 41% and 27% of the *α_max_* for the first, second, and third run, respectively. This confirms that as the organic content increases in the washing system, chlorine loses its efficacy to inactivate microorganisms. This could possibly be due to the competition between various components of organic matter reacting with available free chlorine. As the levels of organic reactants in the wash water increases, chlorine has higher chances of reacting with them and getting depleted, than deactivating *E. coli.* Others (Teng et al., 2018) also have reported that the organic load not only depletes FC but also reduces its sanitizing efficacy. Although the inactivation coefficient was not determined from their studies, a significant decrease in sanitizing efficacy of FC at higher levels of organic load has been reported. In this regard, the efficacy of other sanitizers such as activated persulfate and chlorine dioxide have also been reported to be affected by organic load (Banach et al., 2017; Qi et al., 2020).

**Table 3.** Model parameters for the continuous wash experiments. Errors show the 95% confidence intervals.

| $k_0(\mu M \cdot min^{-1})$ | $\beta_w(MPN \cdot mL^{-1} \cdot min^{-1})$ | $k_M(\mu M)$ | $\beta_{lw}(mL \cdot g^{-1} \cdot min^{-1})$ |
| --- | --- | --- | --- |
| $716 \pm 70.4$ | $45 \pm 1.8$ | $524 \pm 114$ | $0.41 \pm 0.07$ |

Taken together, such an integrated approach in using experiments and mathematical modeling significantly advanced our understanding of the mechanisms by which (i) FC is depleted in single and continuous wash cycles of fresh produce, (ii) FC inactivates pathogens such as *E.coli* in the absence and presence of organic load, (iii) depleting FC levels promote pathogen survival in wash water and their transfer to uncontaminated produce in continuous wash cycles, and finally (iv) determination of model parameters obviates limitations in real-time data collection (e.g., organic load) during continuous produce wash cycles in industrial settings.

#### 4. Conclusions

The current study show that the organic load has a negative effect on the sanitizing efficacy of free chlorine during produce wash, and the chlorine inactivation coefficient for *E. coli* inversely varies with the organic load. Organic load not only consumes free chlorine in produce wash but also decreases chlorine disinfection efficacy, possibly due to competing consumption of FC by the organic matter released from produce cut surface and internal organelles of *E. coli,* although more studies are needed to prove these hypotheses. The disinfection model we developed in this study shows promise in predicting *E. coli* inactivation in the presence of organic load in produce wash processes. Our approach in predicting free chlorine concentration using only a percentage of COD level as an indicator for organic load in continuous wash of iceberg lettuce shows remarkable improvement in the prevailing knowledge of this process. Although the proposed mathematical model was trained on single-wash (batch) experiments, it predicted the free chlorine decay in continuous wash quite accurately. Our proposed model could have significant industrial applications in predicting free chlorine decay during continuous produce wash, usage of minimally appropriate chlorine concentrations to achieve sanitization goals and preventing formation of undesirable disinfection products. Our model, when appropriately calibrated, can predict bacterial survival in wash water as well as help prevent cross-contamination in washing flumes.

## Supplementary Material

### Analysis of variance (ANOVA) for the effect of initial chlorine level and bacterial load on *E. coli* inactivation coefficient

Using the Analysis of Variance, we here show that initial chlorine level (*C_0_*) and bacterial load *(N_0_)* have no effect on the chlorine disinfection efficacy. Since the number of replicates for each combination of factors were balanced in our study, we used Balanced ANOVA. All factors are fixed and so the hypothesis was that all means of inactivation coefficient are equal for different levels of each factor. In other words, *C_0_* and *N_0_* have no effect on chlorine inactivation of *E. coli.* Since the resulting *p*-values were much higher than 0.05 (**Table S1**) from Balanced ANOVA, the difference in mean inactivation coefficients was not significant.

**Table S1.** Balanced Analysis of variance (ANOVA) for the effect initial bacterial level (N_0_) and free chlorine concentration (*C_0_*) on chlorine inactivation coefficient of *E. coli*.

| Source | DF | SS | MS | F | P |
| --- | --- | --- | --- | --- | --- |
| $C_0$ | 2 | 22.59 | 11.30 | 0.23 | 0.794 |
| Log $N_0$ | 1 | 67.25 | 67.25 | 1.40 | 0.257 |
| Error | 14 | 673.84 | 48.13 |  |  |
| Total | 17 | 763.68 |  |  |  |

## Acknowledgements

The authors acknowledge funding from the National Institute of Food and Agriculture Grant No. 2017-67018-26225.

## References

Abnavi, M. D., Alradaan, A., Munther, D., Kothapalli, C. R., & Srinivasan, P. (2019). Modeling of Free Chlorine Consumption and Escherichia coli O157:H7 Cross-Contamination During Fresh-Cut Produce Wash Cycles. Journal of Food Science, 84(10), 2736–2744. doi: 10.1111/1750-3841.14774

Abnavi, M. D., Kothapalli, C. R., & Srinivasan, P. (2021). Total amino acids concentration as a reliable predictor of free chlorine levels in dynamic fresh produce washing process. Food Chemistry, 335, 127651. doi: https://doi.org/10.1016/j.foodchem.2020.127651

Banach, J. L., van Bokhorst-van de Veen, H., van Overbeek, L. S., van der Zouwen, P. S., van der Fels-Klerx, H. J., & Groot, M. N. N. (2017). The efficacy of chemical sanitizers on the reduction of Salmonella Typhimurium and Escherichia coli affected by bacterial cell history and water quality. Food Control, 81, 137–146. doi: https://doi.org/10.1016/j.foodcont.2017.05.044

Bolten, S., Gu, G., Luo, Y., Van Haute, S., Zhou, B., Millner, P.,… Nou, X. (2020). Salmonella inactivation and cross-contamination on cherry and grape tomatoes under simulated wash conditions. Food Microbiology, 87, 103359. doi: https://doi.org/10.1016/j.fm.2019.103359

Chen, X., & Hung, Y.-C. (2017). Effects of organic load, sanitizer pH and initial chlorine concentration of chlorine-based sanitizers on chlorine demand of fresh produce wash waters. Food Control, 77, 96–101.doi: https://doi.org/10.1016/j.foodcont.2017.01.026

Cho M., Kim j., Kim J. Y., Yoon J., Kim J.-H., (2010). Mechanisms of Escherichia coli inactivation by several disinfectants. Water Research, 44, 3410–3418. doi: https://doi.org/10.1016/j.watres.2010.03.017.

Cossu, A., Ercan, D., Wang, Q., Peer, W. A., Nitin, N., & Tikekar, R. V. (2016). Antimicrobial effect of synergistic interaction between UV-A light and gallic acid against Escherichia coli O157:H7 in fresh produce wash water and biofilm. Innovative Food Science & Emerging Technologies, 37, 44–52. doi: https://doi.org/10.1016/j.ifset.2016.07.020

Deborde, M., & von Gunten, U. (2008). Reactions of chlorine with inorganic and organic compounds during water treatment—Kinetics and mechanisms: A critical review. Water Research, 42(1), 13–51. doi: https://doi.org/10.1016/j.watres.2007.07.025

Gibson, K. E., Almeida, G., Jones, S. L., Wright, K., & Lee, J. A. (2019). Inactivation of bacteria on fresh produce by batch wash ozone sanitation. Food Control, 106, 106747. doi: https://doi.org/10.1016/j.foodcont.2019.106747

Gómez-López, V. M., Lannoo, A.-S., Gil, M. I., & Allende, A. (2014). Minimum free chlorine residual level required for the inactivation of Escherichia coli O157:H7 and trihalomethane generation during dynamic washing of fresh-cut spinach. Food Control, 42, 132–138. doi: https://doi.org/10.1016/j.foodcont.2014.01.034

Helbling D. E., VanBriesen J. M. (2007). Free chlorine demand and cell survival of microbial suspensions. Water Research, 41(19), 4424–4434. doi: https://doi.org/10.1016/j.watres.2007.06.006

Hirneisen, K. A., Markland, S. M., & Kniel, K. E. (2011). Ozone Inactivation of Norovirus Surrogates on Fresh Produce. Journal of Food Protection, 74(5), 836–839. doi: 10.4315/0362-028X.JFP-10-438

Jirka, A.M., & Carter, M.J., (1975). Micro-semi-automated analysis of surface and wastewa-ters for chemical oxygen demand. Analytical Chemistry 47, 1397.

Lee, E.-S., Yoon, T.-H., Lee, M.-Y., Han, S.-H., & Ka, J.-O. (2010). Inactivation of environmental mycobacteria by free chlorine and UV. Water Research, 44(5), 1329–1334. doi: https://doi.org/10.1016/j.watres.2009.10.046

Li, J., Teng, Z., Weng, S., Zhou, B., Turner, E. R., Vinyard, B. T., & Luo, Y. (2019). Dynamic changes in the physicochemical properties of fresh-cut produce wash water as impacted by commodity type and processing conditions. PLOS ONE, 14(9), e0222174. doi: 10.1371/journal.pone.0222174

López-Gálvez, F., Truchado, P., Tudela, J. A., Gil, M. I., & Allende, A. (2020). Critical points affecting the microbiological safety of bell peppers washed with peroxyacetic acid in a commercial packinghouse. Food Microbiology, 88, 103409. doi: https://doi.org/10.1016/j.fm.2019.103409

López-Gálvez, F., Tudela, J. A., Allende, A., & Gil, M. I. (2019). Microbial and chemical characterization of commercial washing lines of fresh produce highlights the need for process water control. Innovative Food Science & Emerging Technologies, 51, 211–219. doi: https://doi.org/10.1016/j.ifset.2018.05.002

López-Gálvez, F., Tudela, J. A., Gil, M. I., & Allende, A. (2020). Use of Chlorine Dioxide to Treat Recirculated Process Water in a Commercial Tomato Packinghouse: Microbiological and Chemical Risks. [10.3389/fsufs.2020.00042]. Frontiers in Sustainable Food Systems, 4, 42.

Luo, Y., Nou, X., Millner, P., Zhou, B., Shen, C., Yang, Y.,… Shelton, D. (2012). A pilot plant scale evaluation of a new process aid for enhancing chlorine efficacy against pathogen survival and cross-contamination during produce wash. International Journal of Food Microbiology, 158(2), 133–139. doi: https://doi.org/10.1016/j.ijfoodmicro.2012.07.008

Munther, D., Luo, Y., Wu, J., Magpantay, F. M. G., & Srinivasan, P. (2015). A mathematical model for pathogen cross-contamination dynamics during produce wash. Food Microbiology, 51, 101–107. doi: https://doi.org/10.1016/j.fm.2015.05.010

Qi, H., Wang, L., Huang, Q., & Hung, Y.-C. (2020). Effect of organic load on the efficacy of activated persulfate in inactivating Escherichia coli O157:H7 and the production of halogenated by-products. Food Control, 114, 107218. doi: https://doi.org/10.1016/j.foodcont.2020.107218

Sheng, L., Shen, X., Su, Y., Korany, A., Knueven, C. J., & Zhu, M.-J. (2020). The efficacy of sodium acid sulfate on controlling Listeria monocytogenes on apples in a water system with organic matter. Food Microbiology, 92, 103595. doi: https://doi.org/10.1016/j.fm.2020.103595

Srinivasan, P., Dehghan Abnavi, M., Sulak, A., Kothapalli, C. R., & Munther, D. (2020). Towards enhanced chlorine control: Mathematical modeling for free chlorine kinetics during fresh-cut carrot, cabbage and lettuce washing. Postharvest Biology and Technology, 161, 111092. doi: https://doi.org/10.1016/j.postharvbio.2019.111092

Taylor, R. H., Falkinham, J. O., Norton, C. D., & LeChevallier, M. W. (2000). Chlorine, Chloramine, Chlorine Dioxide, and Ozone Susceptibility of *Mycobacterium avium*. Applied and Environmental Microbiology, 66(4), 1702. doi: 10.1128/AEM.66.4.1702-1705.2000

Teng, Z., Luo, Y., Alborzi, S., Zhou, B., Chen, L., Zhang, J.,… Wang, Q. (2018). Investigation on chlorine-based sanitization under stabilized conditions in the presence of organic load. International Journal of Food Microbiology, 266, 150–157. doi: https://doi.org/10.1016/j.ijfoodmicro.2017.11.027

Tudela, J. A., López-Gálvez, F., Allende, A., & Gil, M. I. (2019). Chlorination management in commercial fresh produce processing lines. Food Control, 106, 106760. doi: https://doi.org/10.1016/j.foodcont.2019.106760

Tudela, J. A., López-Gálvez, F., Allende, A., Hernández, N., Andújar, S., Marín, A.,… Gil, M. I. (2019). Operational limits of sodium hypochlorite for different fresh produce wash water based on microbial inactivation and disinfection by-products (DBPs). Food Control, 104, 300–307. doi: https://doi.org/10.1016/j.foodcont.2019.05.005

Van Haute, S., Luo, Y., Sampers, I., Mei, L., Teng, Z., Zhou, B.,… Millner, P. (2018). Can UV absorbance rapidly estimate the chlorine demand in wash water during fresh-cut produce washing processes? Postharvest Biology and Technology, 142, 19–27. doi: https://doi.org/10.1016/j.postharvbio.2018.02.002

Watson, H. E. (1908). A Note on the Variation of the Rate of Disinfection with Change in the Concentration of the Disinfectant. The Journal of hygiene, 5(4), 536–542. doi: 10.1017/s0022172400015928

